# Wrack enhancement of post-hurricane vegetation and geomorphological recovery in a coastal dune

**DOI:** 10.1101/2021.03.12.435051

**Authors:** Matthew A. Joyce, Sinead M. Crotty, Christine Angelini, Orlando Cordero, Collin Ortals, Davide de Battisti, John N Griffin

**Affiliations:** Department of Biosciences, Swansea University, Swansea, UK; Department of Environmental Engineering Sciences, University of Florida, Gainesville, Florida, US; Carbon Containment Lab, Yale School of the Environment, Yale University, New Haven, Connecticut, US; Department of Civil and Coastal Engineering, University of Florida, Gainesville, Florida, US; Department of Geological Sciences, University of Florida, Gainesville, Florida, US; Department of Biology, University of Pisa, Pisa, Italy

**Keywords:** Coastal resilience, wrack, marine subsidies, coastal sand dunes, ecosystem recovery, hurricane

## Abstract

Coastal ecosystems such as sand dunes, mangrove forests, and salt marshes provide natural storm protection for vulnerable shorelines. At the same time, storms erode and redistribute biological materials among coastal systems via wrack. Yet how such cross-ecosystem subsidies affect post-storm recovery is not well understood. Here, we report an experimental investigation into the effect of storm wrack on eco-geomorphological recovery of a coastal embryo dune in north-eastern Florida, USA, following hurricane Irma. We contrasted replicated 100-m^2^ wrack-removal and unmanipulated (control) plots, measuring vegetation and geomorphological responses over 21 months. Relative to controls, grass cover was reduced 4-fold where diverse storm wrack, including seagrass rhizomes, seaweed, and wood, was removed. Wrack removal was also associated with a reduction in mean elevation, which persisted until the end of the experiment when removal plots had a 14% lower mean elevation compared to control plots. These results suggest that subsides of wrack re-distributed from other ecosystem types (e.g. seagrasses, macroalgae, uplands): i) enhances the growth of certain dune-building grasses; and ii) boosts the geomorphological recovery of coastal dunes. Our study also indicates that the practice of post-storm beach cleaning to remove wrack – a practice widespread outside of protected areas – may undermine the resilience of coastal dunes and their services.

## INTRODUCTION

Anthropogenic climate change is disrupting the physical and biological structure of ecosystems worldwide, threatening the continued delivery of ecosystem services and human wellbeing (Mooney *et al*., 2009; Nelson *et al*., 2013; Scholes 2016). Increasing climatic variability and extreme weather events such as droughts, heat waves, floods and hurricanes are emerging as key signatures of anthropogenic climate change (Pohl *et al*., 2017; Ummenhofer & Meehl 2017), and are amplifying perturbations to ecosystems (Jentsch & Beierkuhnlein 2008; Altwegg *et al*., 2017). Identifying controls on the capacity of ecosystems and their services to recover from extreme events is a major research challenge.

Evidence from multiple ecosystems indicates that connectivity and cross ecosystem subsidies can support recovery following massive disturbances. Connectivity supplements many systems with trophic resources, recruits, and facilitative species (Moore *et al*., 2007; Adam *et al*., 2011; Mueller *et al*., 2014). Such cross-ecosystem connections have been observed to enhance ecosystem recovery from major disturbances including forest fires (Cavallero *et al*., 2013), droughts (Sarremejane *et al*., 2020), and mass coral bleaching (Adam *et al*., 2011; Hock *et al*., 2017). Cross-system subsidies are expected to be especially important in supporting ecosystem recovery in typically unproductive environments such as deserts, the deep sea, and coastal dunes (Spiller *et al*., 2010; Schlacher *et al*., 2013; Filbee-Dexter *et al*., 2018), as well as in cases where such materials stabilize substrates otherwise vulnerable to erosion or rapid geomorphic change.

Coastal dunes occupy over 30% of the world’s ice-free coastlines and in many regions provide a natural flood defence against storm surges and waves which pose a threat to life, infrastructure, ecosystems and agriculture (Sherman-Morris *et al*., 2015). Plants – particularly grasses – often play an important role in dune formation, as they capture wind-blown sediments which in turn stimulates their growth (Miller *et al*., 2009). Following storms cross ecosystem detrital inputs delivered by tides (‘wrack’) are likely to capture sediment particles and fuel growth of ecosystem engineering plants, promoting dune recovery. As large storms and hurricanes cause widespread erosion in coastal ecosystems, such as seaweed and seagrass beds, mangroves and saltmarshes (Stone *et al*., 1997), large and compositionally diverse wrack is likely to be deposited after major storms. Indeed, previous studies have identified the potential of wrack (predominantly seaweed) to stimulate dune plant growth by providing a nutrient source (Williams & Feagin 2010; Dufour *et al*., 2012; MacMillan & Quijón 2012). However, estimates of how realized deposits of storm wrack, which are likely to be increasingly large and diverse, influence post-storm vegetation and geomorphological recovery are currently lacking. Understanding the role of storm wrack in dune recovery has applied relevance because storm wrack and other storm debris are often removed by managers in a bid to improve public beach safety and aesthetics (Gilburn 2012). Here, we report an experimental test of storm wrack effects on dune vegetation and geomorphological recovery following Hurricane Irma in September of 2017. Storm surges associated with this disturbance event caused widespread dune erosion along Florida’s coastline (Florida Department of Environmental Protection 2018; Nagarajan *et al*., 2019) and a substantial deposition of wrack material in these areas. To quantify the role of cross-system wrack subsidy on coastal ecosystem recovery trajectories, we conducted a large-scale wrack removal experiment at a site in north-eastern Florida that is protected from beach cleaning activities common in the region. We hypothesized that: i) cross-system wrack subsidy enhances plant abundance and recovery, especially that of dune-building grasses; and ii) wrack presence facilitates the accretion of sediment and overall elevation gain of re-forming embryo dunes.

## METHODS AND MATERIALS

The study was conducted at Anastasia State Park (hereafter ‘ASP’), Florida, US (29°87’75” N, 81°27’13” W). ASP is a state managed park, containing a continuous coastal sand dune system approximately 6 km long, sparsely punctuated by dune blow-outs, reaching 7.7 m above mean sea level (datum: NAD 1983) in height. Fore- and embryo-dunes in the region of the study site - characteristic of dunes in temperate and subtropical zones worldwide - contain mixed communities of dune-building grasses, forbs, and vines (Overlease 1991). In foredunes *Uniola paniculata* (sea oats) is the dominant grass and a key dune builder, while another dune builder *Panicum amarum* (panic grass) and the dune stabiliser *Sporobolus virginicus* (seashore dropseed) are also common. Dunes in the region sit within a diverse suite of coastal ecosystems, located within a latitudinal mangrove – saltmarsh transition zone, and with both nearby floating seaweed (*Sargassum*) and - further south - seagrass meadows.

We conducted our experiment in a 1.3 km long swathe of storm wrack deposited in the supratidal zone at ASP. Storm wrack formed a continuous band in the embryo dune and was comprised of plant, macroalgae and anthropogenic materials. This zone was 10 – 12 m wide and up to 60 cm deep. On our first visit to the site in early February 2018 we determined this wrack zone to be associated with a recent hurricane (hurricane Irma) as: i) there was an abnormal amount of wrack deposited in the supratidal (which can only occur during storm surge events) and ii) the wrack contained a variety of organic matter typical of heavily eroded systems, including seagrass rhizomes. These assumptions were confirmed by liaising with land managers at ASP. To set up the experiment, five blocks were distributed along 1.3km within the storm wrack zone (see Supplementary Materials, Fig. S1), with each block containing three, 10 x 10 m plots: a wrack removal plot, a procedural control plot and a control plot. Plots were spaced at least ten meters apart in randomized orders (along the beach) in each block; blocks were spaced at least 80 meters apart, 0.7 - 2 km from public beach entrances. In wrack removal plots, care was taken to avoid disturbing vegetation by removing wrack in the vicinity of plants carefully using hand tools. Storm wrack was extracted up to a depth of 60 cm by sieving through 0.5 cm wide mesh openings over removal plots. Procedural controls were treated using gardening forks to lift wrack buried up to 30 cm deep, before placing it back in its original position to simulate removal plot disturbance (Supplementary Materials, Fig. S2). We utilized a disturbance depth of 30 cm as opposed to 60 cm in procedural control plots as both wrack material and potentially disturbed plant roots were concentrated in this upper zone. Control plots remained undisturbed. Experimental set-up was conducted between the end of March and early May 2018.

### Data collection

Following experiment set-up, a grid was marked out in each plot, producing nine data collection points at bisecting points of lateral orthogonal transects three, five and seven meters along on plot axes (Supplementary Materials, Fig. S3). Small (0.25 m^2^) quadrats at each grid point were used to estimate vegetation composition and abundance immediately following deployment on early May 2018, early August 2018, mid-October 2018, late January 2019, and finally in July 2019. We use all time points to describe changes in dune grass abundance in controls but focus on the final time point in comparisons to best represent the long-term consequences of wrack removal. We collected geomorphological data upon deployment and on three occasions following the completion of the experiment to both characterise the overall elevation of the control and removal plots, and to explore the impact and recovery in the face of a subsequent tropical storm. This data was collected prior to tropical storm Dorian in late August 2019, immediately before the storm in early September 2019 and once more in December 2019. Geomorphological data was collected using a Riegl VZ-400 3D laser scanner which produced 3D computational digital elevation models of each experimental plot. Post-processing using RiSCAN Pro allowed for the calculation of the mean elevation above sea level (m). All elevations measured (and referred to hereafter) are based on the Datum NAD 1983 (Conus) in which mean sea level is absolute zero elevation. Storm Dorian passed the study site at mid-tide and therefore only had minor and accretional impacts on most of the plots. Accordingly, our main analysis focuses on the first and final timepoint (May 2018 and December 2019) to illustrate the total accumulated difference in geomorphology between treatments. A breakdown of storm impact and recovery metrics is presented in the supplementary materials (Fig. S8).

To characterise storm wrack composition in the study plots, all wrack extracted from removal plots was collected into heavy duty refuse sacks. Between three to four sacks were randomly selected from each block for detailed analysis. Wrack was separated into component parts, which were then dried and weighed. C:N analysis was conducted on each organic wrack type using a Carlo Erba NA1500 CNHS elemental analyzer. For C: N, three samples per wrack type per experimental replicate were analyzed. To additionally characterise the composition of wrack in the broader region following hurricane Irma (in February 2018), we surveyed wrack composition across 21 sites along 200km of coastline in Florida from Cape Canaveral National Seashore (28°55’42.72” N, 80°49’20.19” W) to Fernandina Beach (29°52’29.92” N, 81°16’11.90” W). The surveys were conducted as part of a companion project and therefore included a range of landscape development contexts. The mean cover of wrack types were estimated visually along three, 30m alongshore transects in the (storm) wrack zone at each site.

### Data analysis

All analyses were run in R version 3.4.1 (R Core Team, 2020). To test the effects of treatment on ecological and geomorphological response variables, we used linear models focusing on the first and final time point for geomorphological responses (May 2018 and December 2019) and final timepoint for vegetation (July 2019) responses, including treatment and block as additive fixed factors. Block was treated as a fixed (rather than random) factor as it allows for a simple and compact summary of differences between treatments (Dixon 2016). All results were qualitatively identical whether block was considered fixed or random. Initial analyses showed that controls and procedural controls were indistinguishable, allowing us to pool data for these treatments to increase statistical power when contrasting to the removal treatment. All plant cover variables were log (+1) transformed prior to analysis to improve residual distributions.

## RESULTS

### Wrack composition

Wrack collected from removal treatment plots consisted of seven main components (Fig. 1). The most common components by mass were rhizomes from the seagrass *Thalassia testudinum* (44%), followed by vascular plant material consisting mostly of the salt marsh cordgrass *Spartina alterniflora* (19%) (Fig. 1A). Pieces of wood were the next most abundant (19%), followed by uprooted *U. paniculata* (7%), mangrove seeds (4%), plastic or other anthropogenic materials (4%) and *Sargassum sp*. (3%). Relative abundances of wrack types were largely consistent across experimental blocks, with seagrass rhizomes consistently the largest component and vascular plant material and wood consistently in the top three (Supplementary Materials Fig. S4). Across the 21 surveyed sites in north-eastern Florida, wrack composition was broadly similar to that at ASP, with seagrass rhizomes and vascular plants the main components (Fig. 1B). Wrack components collected from removal plots at ASP varied widely in C:N ratio, from relatively low-quality wood and uprooted sea oats (*U. paniculata*) to the relatively high nutrient quality mangrove seeds and *Sargassum* (Fig. 1C).

**Figure 1:**
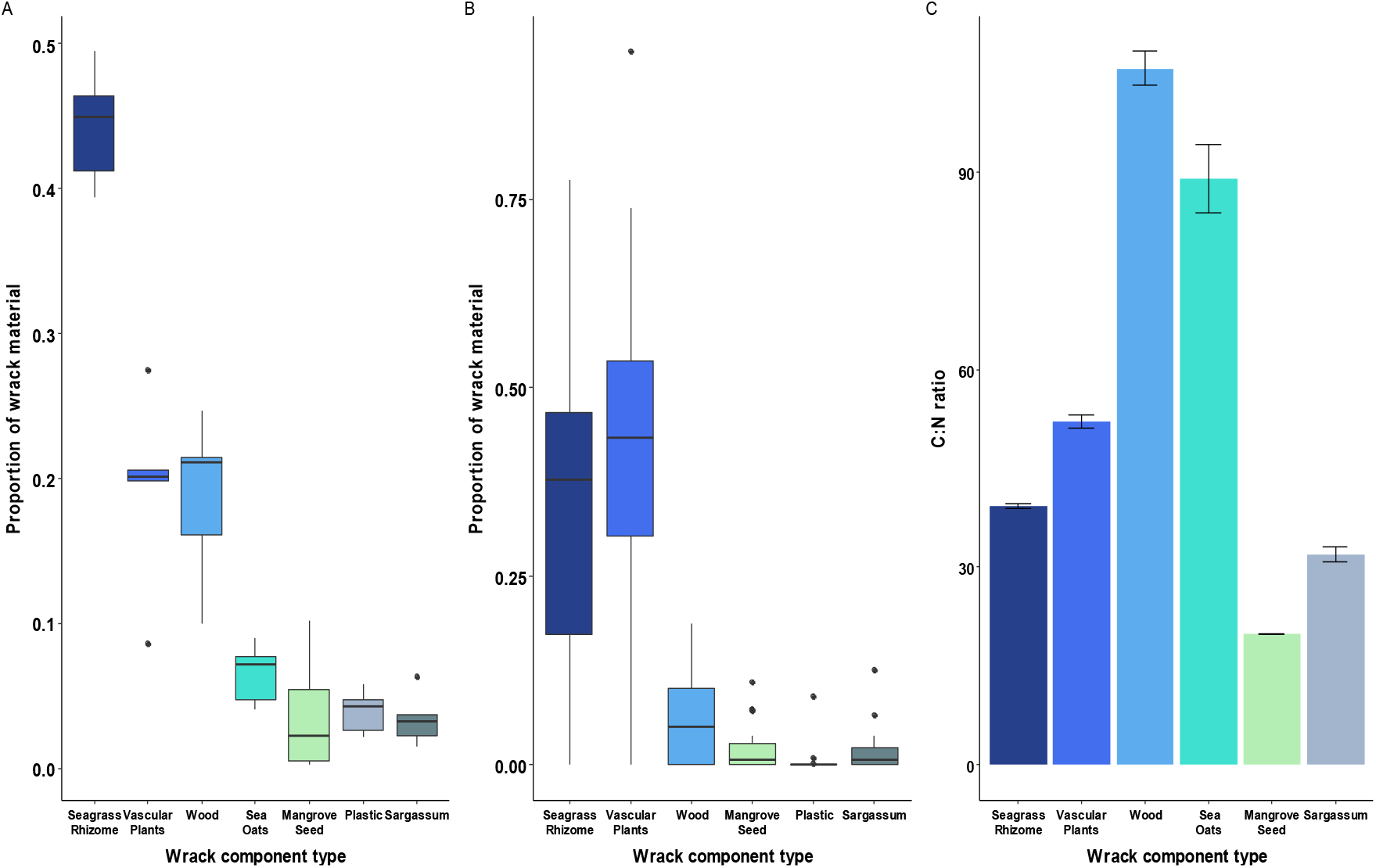
The composition of storm wrack in experimental plots and eastern Florida coastline. Proportions of different components of storm wrack **A)** collected from experimental removal plots (see Supplementary Materials Fig. S4 for proportions in each block) and **B)** from surveys of wrack observed at 21 coastal dune sites along the north-east Florida coastline following hurricane Irma. **C)** Mean (± standard error; n = 15) C:N ratio of organic materials from experimental wrack removal plots.

### Removal effects on vegetation and geomorphology

We first examine the response of vegetation to wrack removal in our embryo dune experimental plots. In control plots, the cover of all plants, including the three grass species (*U. paniculata, P. amarum and S. virginicus*) increased during the experiment (Fig. 2). Of these species, *P. amarum* (final cover: 10.9%) increased substantially, *S. virginicus* (4.5%) moderately, and *U. paniculata* minimally (0.3%). Compared to control plots, storm wrack removal decreased the overall cover of plants (Fig. 3A; t = -2.273, p = 0.049) and the overall cover of grass species (Fig. 3B; t = -5.585, p < 0.001). Storm wrack removal strongly and consistently decreased the cover of *P. amarum* (Fig. 3C; t = -10.885, p < 0.001). Wrack removal also reduced the cover of *S. virginicus* (Fig. 3D; t = -2.855, p = 0.019). Although *U. paniculata* achieved very low abundance in both controls and removals, its cover was increased by wrack removal (Fig. 3E; t = 3.518, p = 0.007) suggesting the conditions within the storm wrack did not favour this species. By the end of the experiment there was a mean of 4.67 plant species per plot, with no detectable differences in species richness between control and removal plots (see Supplementary Materials Fig. S5).

**Figure 2:**
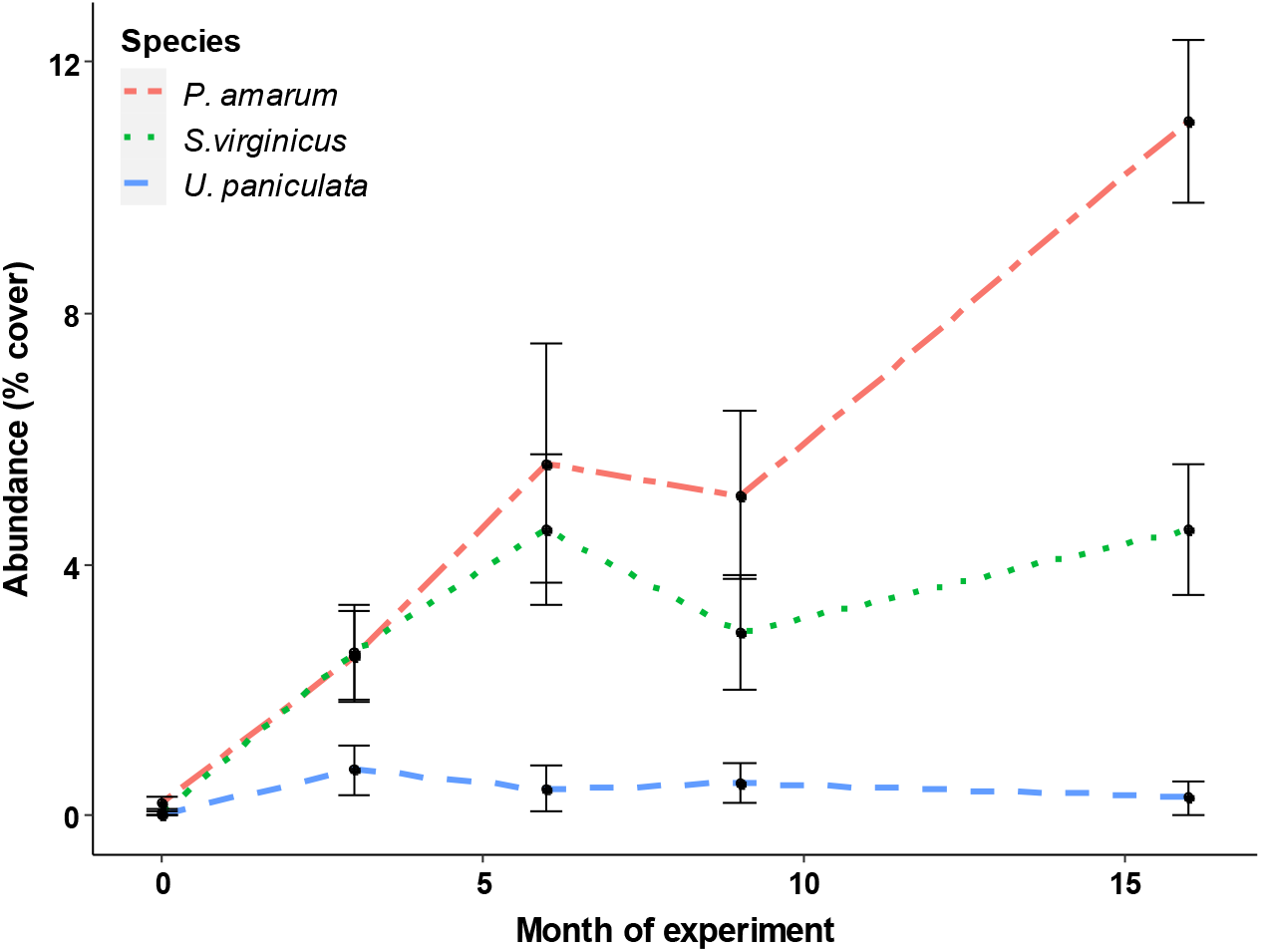
Changes in the abundance of grass species in control plots during the first 16 months of the experiment. Mean (± se) cover of *P. amarum, S. virginicus* and *U. paniculata* in control plots (March 2018 – July 2019).

**Figure 3:**
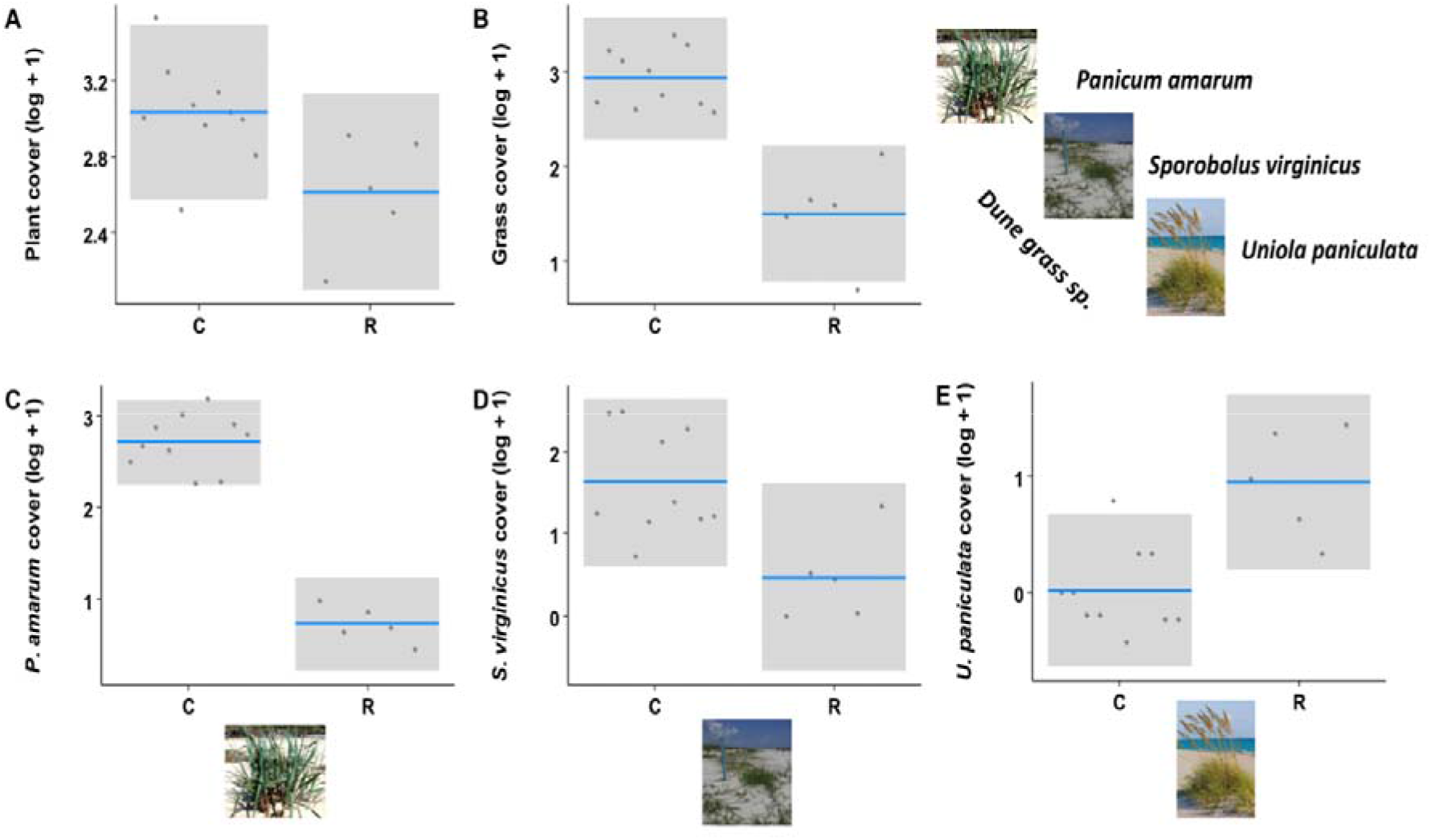
Effects of wrack removal on plant variables. Control is denoted by C and removal by R. All plots show the partial residuals from linear models accounting for experimental block as a function of treatment type. The variables shown are: log-transformed cover of all plants **(A)**, all grasses **(B)**, *P. amarum* **(C)**, *S. virginicus* **(D)**, and *U. paniculata* **(E)**. Plant variables were taken after 16 months. Grey area are the 95% confidence intervals of the means.

Geomorphological responses also differed between control and wrack removal plots (see Fig. 4A and B for visual representation of example plot elevations). Shortly after experimental deployment, there was no clear effect of wrack removal on mean elevation (Fig. 4C; t = -1.34, p = 0.213). After 21 months of monitoring, wrack removal plots had 14% lower mean elevation than controls (Fig. 4D; t = -2.57, p = 0.033). This effect was also qualitatively consistent when considering mean elevation over the final three sets of measurements (see Supplementary Materials Fig. S6). Despite the observed differences in mean elevation at the end of the experiment, there was no significant effect of wrack on the growth of dunes during the experiment, i.e., the difference between initial and final mean elevations (Fig 4E; t = -0.641, p = 0.538). Additionally, there were no clear differences between treatments immediately after or in the period following tropical storm Dorian (Supplementary Materials, Fig. S7), suggesting wrack presence did not play a significant role in accreting material or preventing erosion of sand dunes in this period.

**Figure 4:**
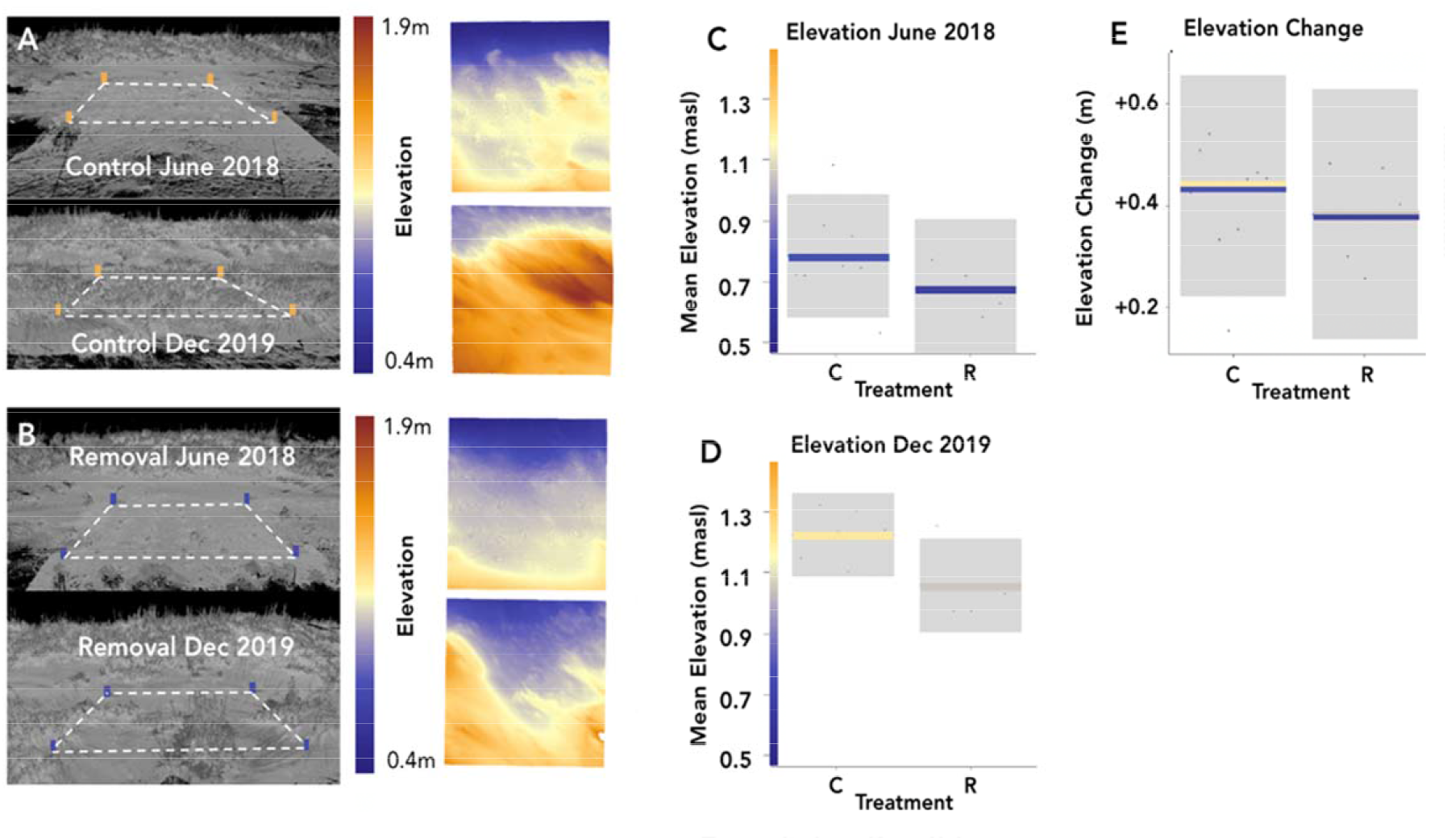
Effects of wrack on geomorphological variables. Laser scan images of control **(A)** and removal **(B)** plots from initial and final geomorphological data collection time points and respective heat maps showing changes in elevation between the two time points. Blue edge to heat map represents seaward side of dune. **C-E:** All plots show the partial residuals from linear models accounting for experimental block as a function of treatment type. Control is denoted by C and removal by R. The variables shown are: mean elevation above sea level at initial data collection point **(C)** and final data collection point **(D)**, and difference of elevation between first and final time point (elevational change) **(E)**. Grey area are the 95% confidence intervals of the mean.

## DISCUSSION

Our study provides evidence that storm wrack enhances vegetation and increases coastal dune elevation following hurricanes. To our knowledge this is the first experiment to investigate the role of wrack associated with a tropical storm on the recovery of a dune plant community. Experimental plots which retained storm wrack showed a greater increase in the cover of several grass species and had a higher elevation and at the end of the experiment. These findings show that where possible, coastal managers should retain storm wrack to aid natural dune recovery.

Wrack enhanced the overall vegetation recovery and had particularly strong effects on several grass species. Storm wrack likely boosted plant establishment and growth by provisioning limiting nutrients. Indeed, macrophyte wrack has been previously shown to enhance N content of sandy interstitial spaces (Dugan *et al*., 2011), while NPK or seaweed supplementation has been shown to elicit positive growth responses in dune building grasses (Hester & Mendelssohn 1990; Williams & Feagin 2010). Notably, while previous studies have focused on single wrack types, we found that storm wrack along Florida’s north-eastern coastline and at our focal study site was highly diverse, representing inputs from a range of coastal ecosystems including seagrass meadows, saltmarshes, mangrove forests, floating seaweed beds, and dunes themselves. Particularly notable was the large component of seagrass rhizomes, particularly at our study site. The large volume indicates these materials may have provided a substantial boost to plant growth in the experiment. The nearest mapped seagrass habitats are ∼100km south at Cape Canaveral, suggesting these materials had travelled considerable distance after storm disturbance or that seagrass was transported from closer, unmapped, beds. Despite being globally threatened (James *et al*., 2020; Strydom *et al*., 2020), seagrass habitats along the Florida coastline have benefited from restorative efforts in recent years (Rezek *et al*., 2019). Our results emphasise the importance of the continued conservation of these ecosystems and attendant cross-system subsidies in Florida. Vascular plant material (comprised mostly of the salt marsh grass *S. alterniflora*) was another major component of storm wrack. Accordingly, as latitudinal shifts of mangroves displace salt marshes along the south-east US coast (Cavanaugh *et al*., 2019), coastal dunes may experience a reduction of valuable saltmarsh detritus. The diverse wrack components exhibited broad variation in carbon to nitrogen ratio, which may ensure both short- and long-term nutrient provision (Enríquez *et al*., 1993). We hypothesise that the high quality *Sargassum* wrack that is quick to decompose provided an early initial pulse of nutrients, while more carbon rich components such as wood will likely provide a longer-term nutrient source. Wrack may also have enhanced plant cover via other mechanisms such as trapping wind-blown seeds (Dugan & Hubbard 2010), increasing water retention, and modulating environmental extremes, including high temperatures (Colombini *et al*., 2000).

Our results also show that wrack can change the relative abundance of several dune grass species. Particularly notable was the strong positive response of *P. amarum* to wrack, contrasted by the negative response of *U. paniculata* (which remained at low abundance in both treatments). These differences may partly reflect interspecific differences in tolerance to salinity and salt spray found at the relatively low elevations of the storm wrack zone, with *P. amarum* more tolerant to these stressors than *U. paniculata* (Dahl & Woodward 1977; Lonard *et al*., 2011). Further, the presence of low-quality materials such as wood may eventually generate humic soils to which *P. amarum* is relatively better suited (Bachman & Whitwell 1995; Willis & Hester 2008). Finally, the colonisation rate of these species differs, with *P. amarum* quickly recolonising dunes (Lonard & Judd 2011), whist *U. paniculata* may take multiple seasons to become established (Lonard *et al*., 2011). As *U. paniculata* is a key dune building species, its response to storm wrack warrants further investigation including studies that evaluate dune succession over time periods longer than our study. More generally, the extent to which dune grass species differentially respond to storm wrack deserves further work as it may have long term consequences for dune geomorphology and services (Reijers *et al*., 2019, Hacker *et al*., 2011).

Our finding that wrack was associated with a greater dune elevation after 21 months of the experiment illustrates that wrack (and its removal) can have persistent impacts on the geomorphology of embryo dunes. However, we detected no clear effect of wrack on changes in elevation - thus dune growth - during the experiment. This suggests that differences were largely established at the start of the experiment and persisted – but were not further amplified - during geomorphological recovery. The lack of an effect of dune growth is surprising considering the clear influence of wrack removal on overall grass cover and specifically *P. amarum* cover, which would be expected to influence capture of sediment as seen in previous restorative efforts (Dahl & Woodward 1977). It is possible that negative wrack removal effects on *P. amarum* were offset by positive effects on *U. paniculata*, although this is unlikely considering the low abundance of *U. paniculata*. Alternatively, it is possible that dune growth is more strongly determined by aeolian processes rather than interactions with plants in the early stages of dune recovery (and therefore relatively low cover of dune-building grasses) in this system. Notwithstanding similar dune growth in wrack presence and absence plots, wrack removal had a persistent negative effect on dune geomorphology, suggesting that, in addition to the previously discussed impact on plant communities, beach management practices such as the mechanical raking of wrack (Nordstrom *et al*., 2011) are likely to reduce dune elevation during storm recovery.

In summary, our results build on previous studies which have investigated the effects of small-scale wrack additions on undisturbed dunes (Hooton *et al*., 2014), by showing novel direct experimental evidence that removal of storm wrack hinders vegetation development. In addition, we show that while not contributing to overall dune growth rate, wrack removal reduced overall elevational gains. Considering these findings, the common management practise of wrack and debris removal following major disturbances should be revised to allow the retention of beneficial organic materials. Our results illustrate that not only do storms erode coastal ecosystems, but redistribution of liberated materials can support resilience in recipient systems. Given the risk of increased storm intensity and frequency in coastal areas (Wang & Toumi 2021), compounded by accelerating sea level rise (Horton *et al*., 2020), further understanding – and sensitive management – of natural processes supporting coastal ecosystem resilience must be a priority.

## ACKNOWLEDGMENTS

This work was funded by a NERC Urgency grant to JG (NE/R016593/1) and an NSF CAREER Grant to CA (1652628). We thank Alice Bard Florida Department of Environmental Protection, and Mark Giblin and Jason Dupoy at the Anastasia State Park for facilitating our research. We also thank T. Fairchild, S. Sharp, and A. Bersoza for valuable assistance in the field.

## SUPPLEMENTARY MATERIALS

**Fig. S1:**
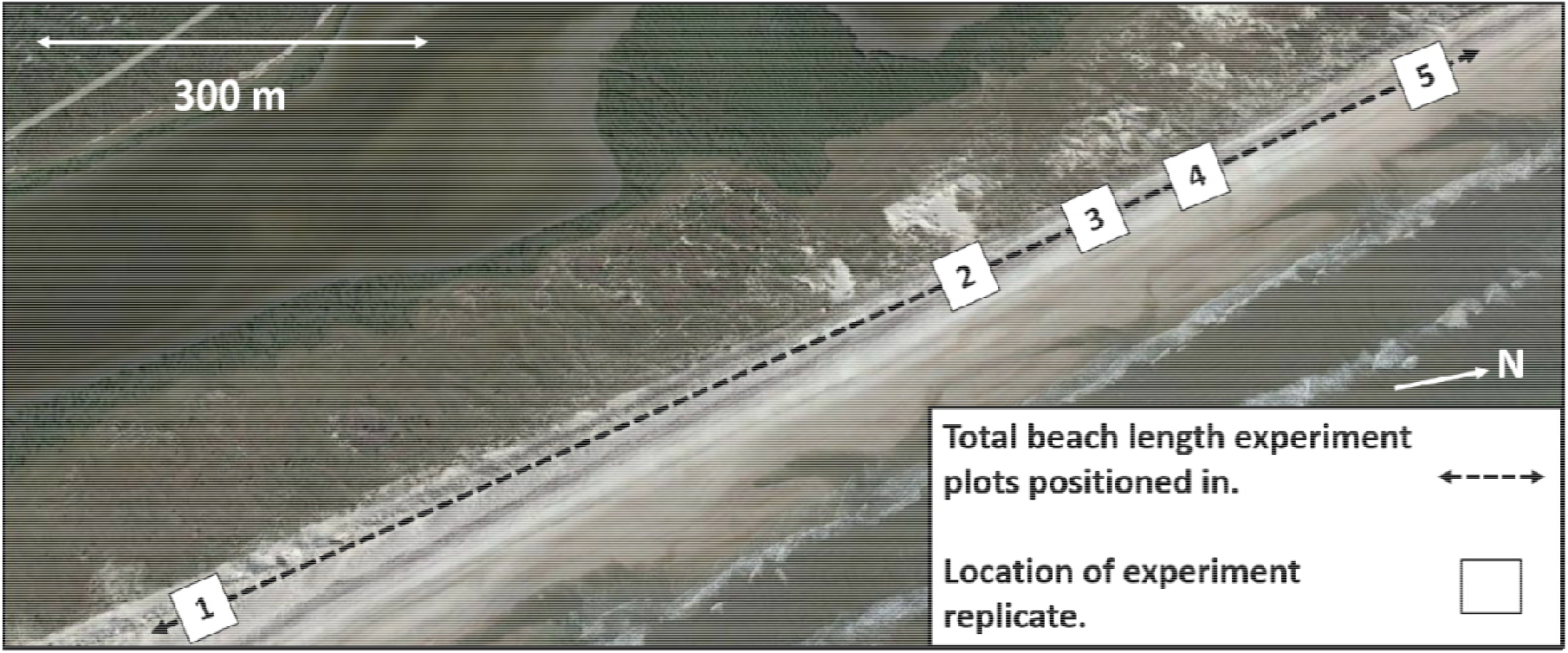
Google Earth composite image of Anastasia experimental site (30/10/18), illustrating the location of blocks (1 – 5) on sand dune system.

**Fig. S2:**
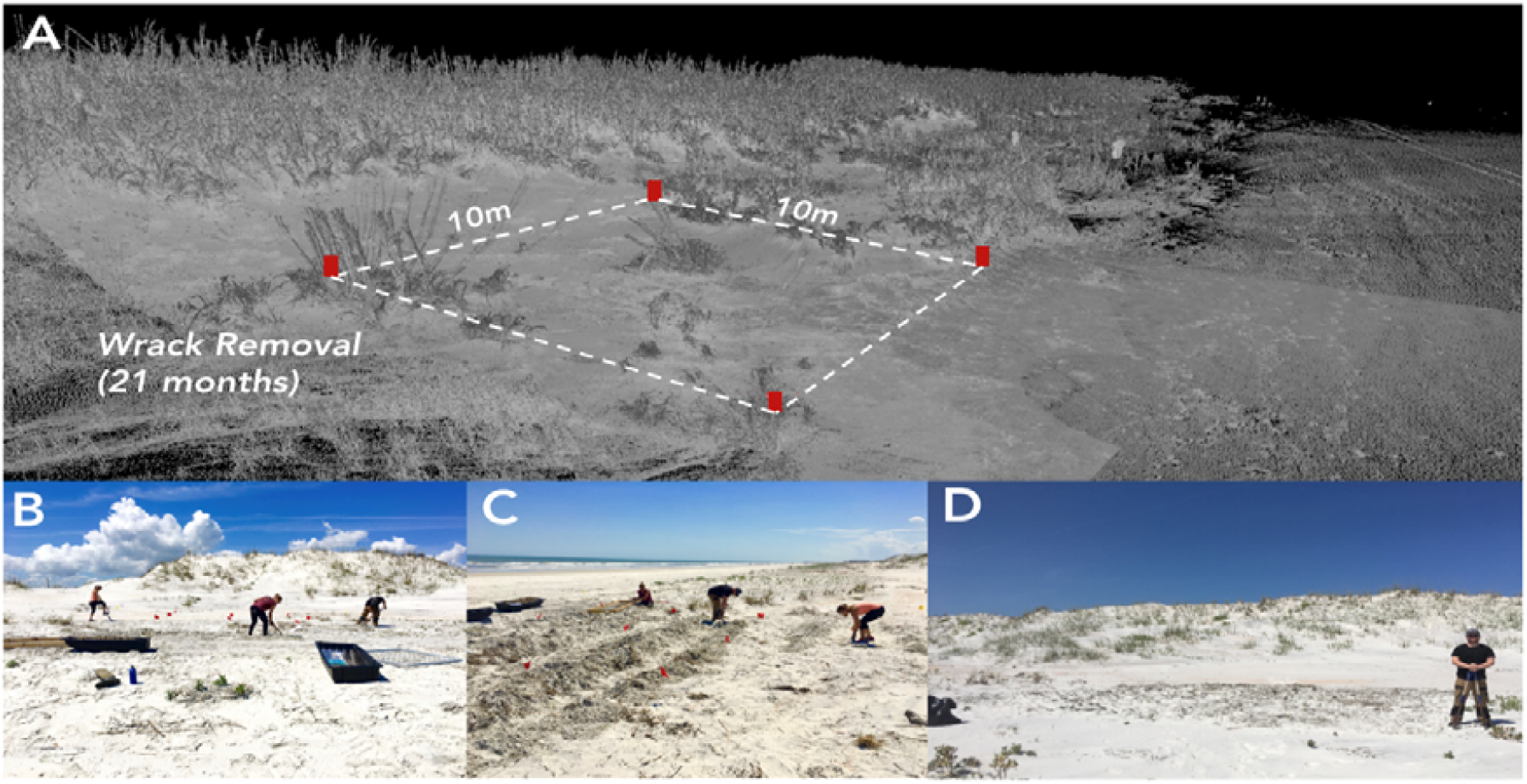
Laser scan of removal plot and photographs of deployment of typical 10 x 10 m experimental plot in storm wrack zone at Anastasia. **A)** Laser scan image shows removal plot 21 months after experimental deployment, red blocks represent corners of 10 x 10 m plot. **B – C)** Photographs of surveyors digging wrack out of removal plot prior to sieving process during experiment deployment. **D)** Photograph showing removal plot (bare area of sand to rear left of surveyor) immediately after deployment completed.

**Fig. S3:**
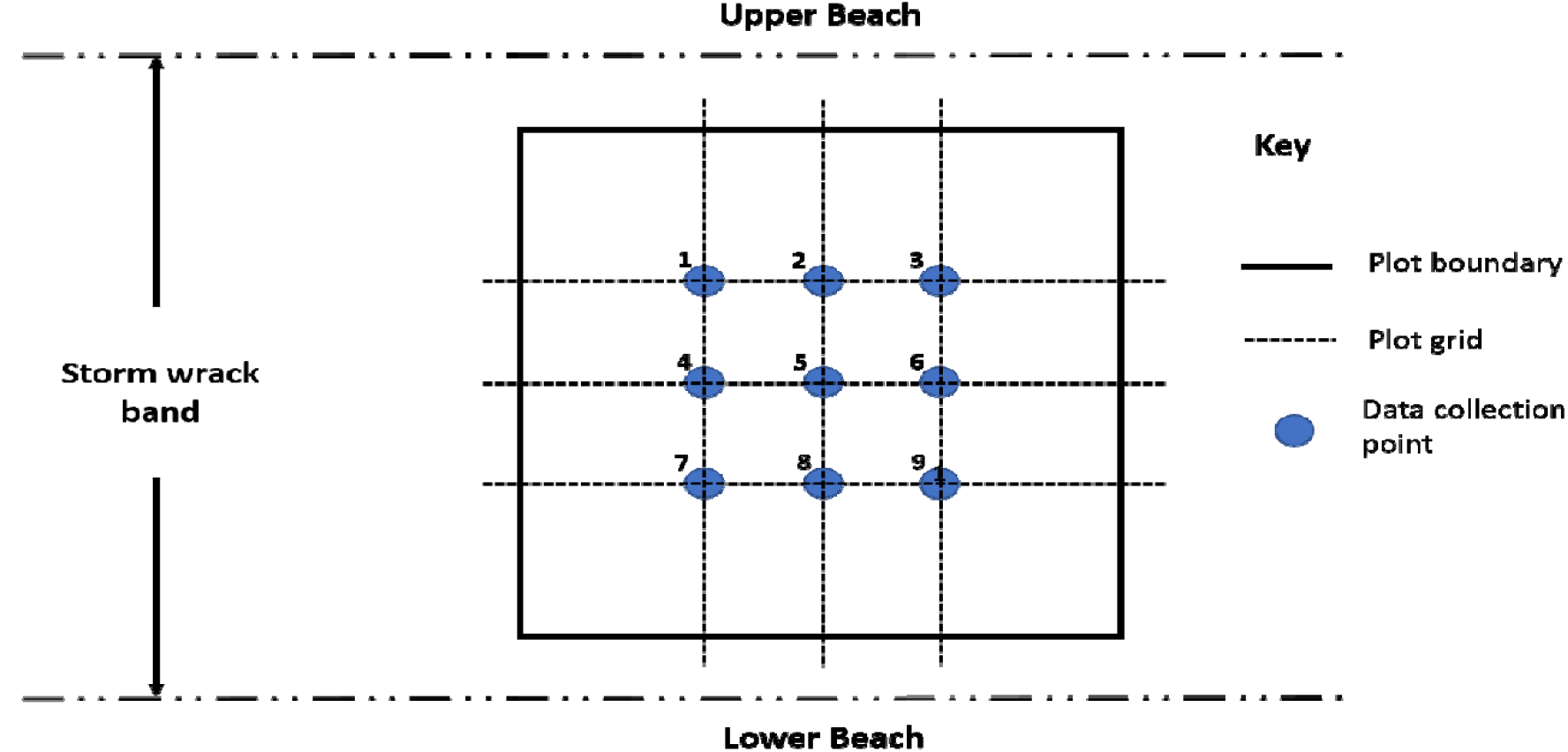
Anastasia experimental plot schematic. Schematic shows nine data collection points at bisecting points of lateral orthogonal transects three, five and seven metres along on plot axes. Data collection points show positions where 0.25 m^2^ quadrats were used to estimate vegetation abundance.

**Fig. S4:**
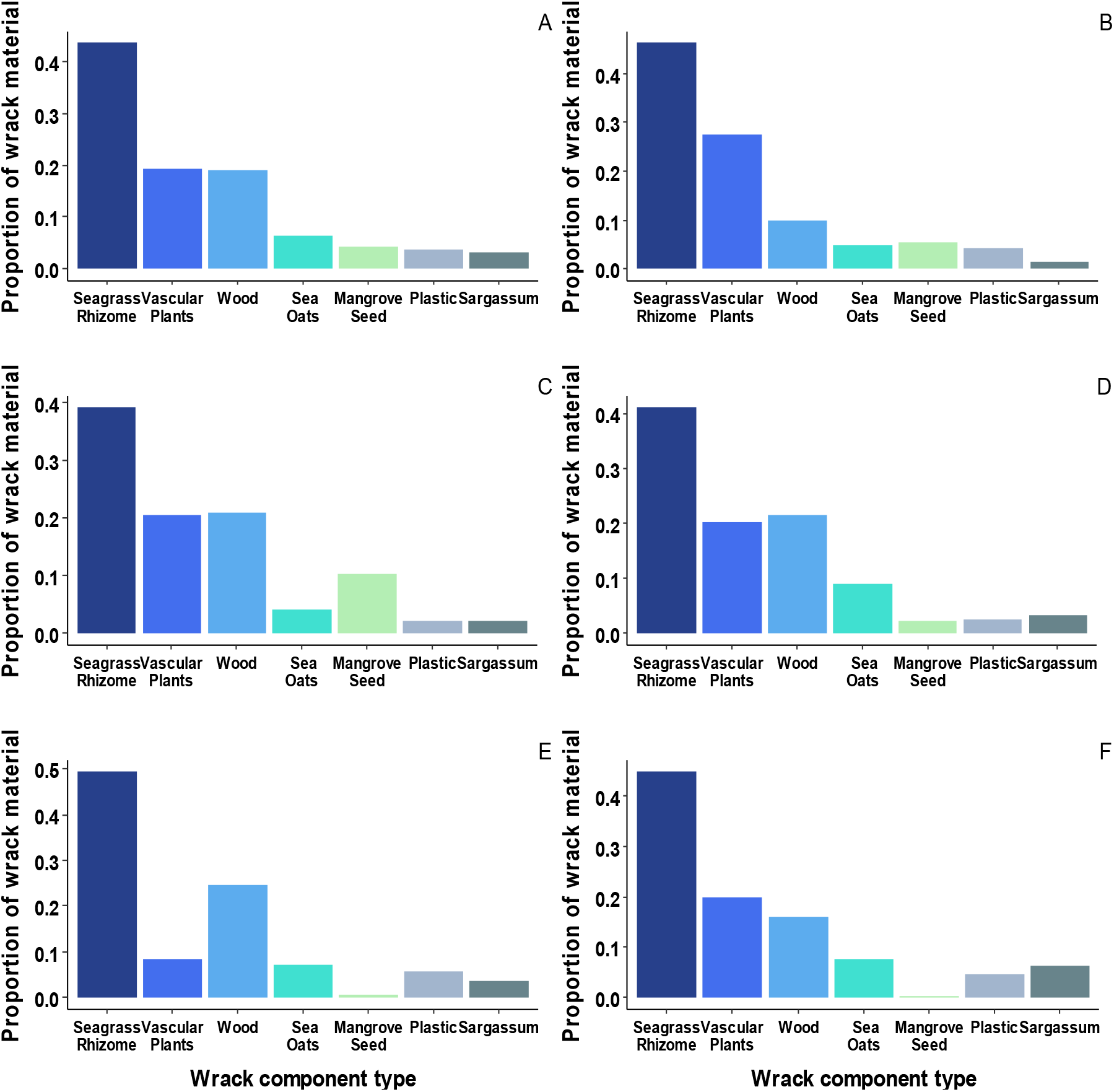
The proportional composition of storm wrack in experimental plots. Fig. shows the proportions of different components of storm wrack removed from all blocks combined (A) and from individual experimental plots (B-F).

**Fig. S5:**
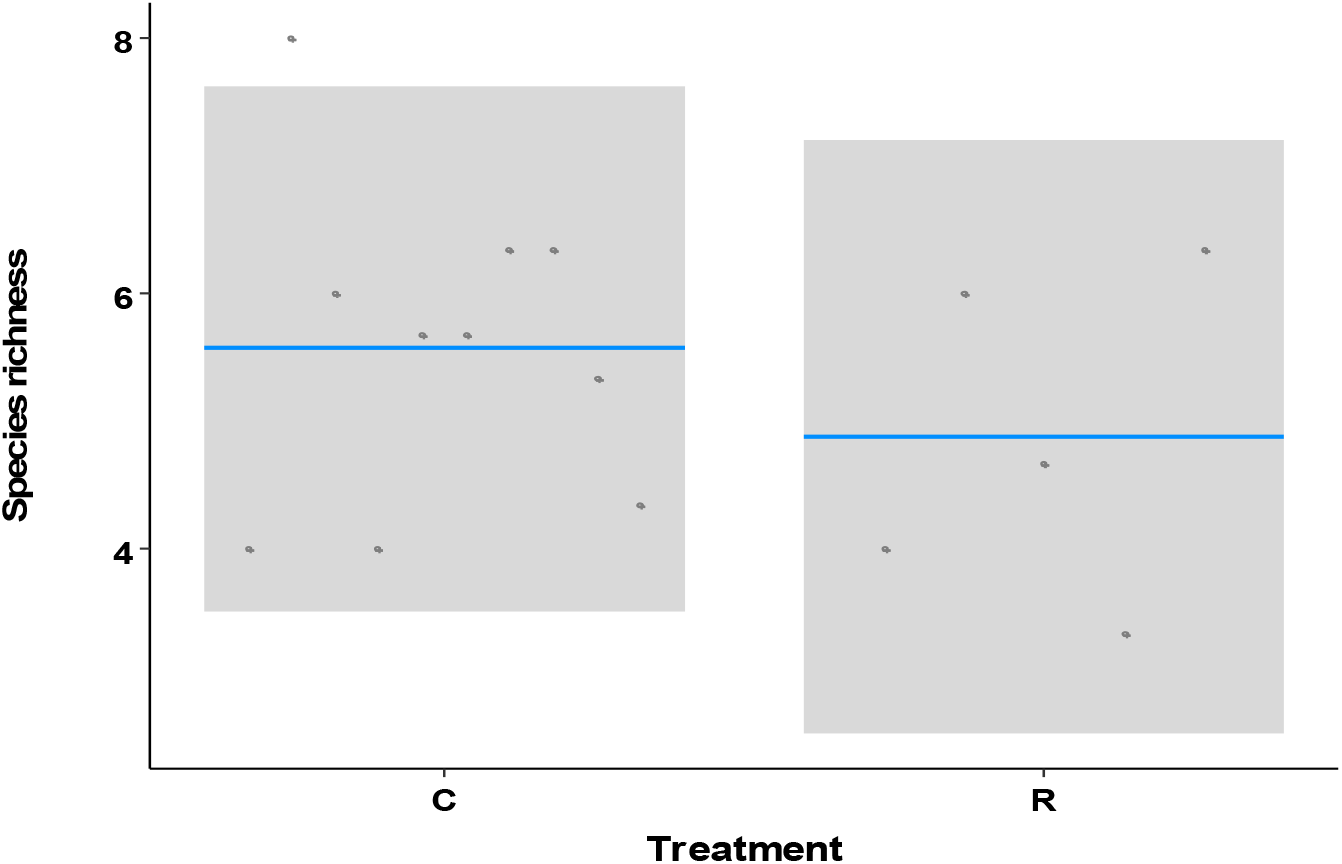
Effects of wrack on plant species richness. Plot shows the partial residuals of linear model showing the predicted species richness as a function of treatment type (control vs removal). Measurements taken after 15 months. No significant difference detected (t = - 0.850, p = 0.42). Grey area represents the 95% confidence intervals of the means.

**Fig. S6:**
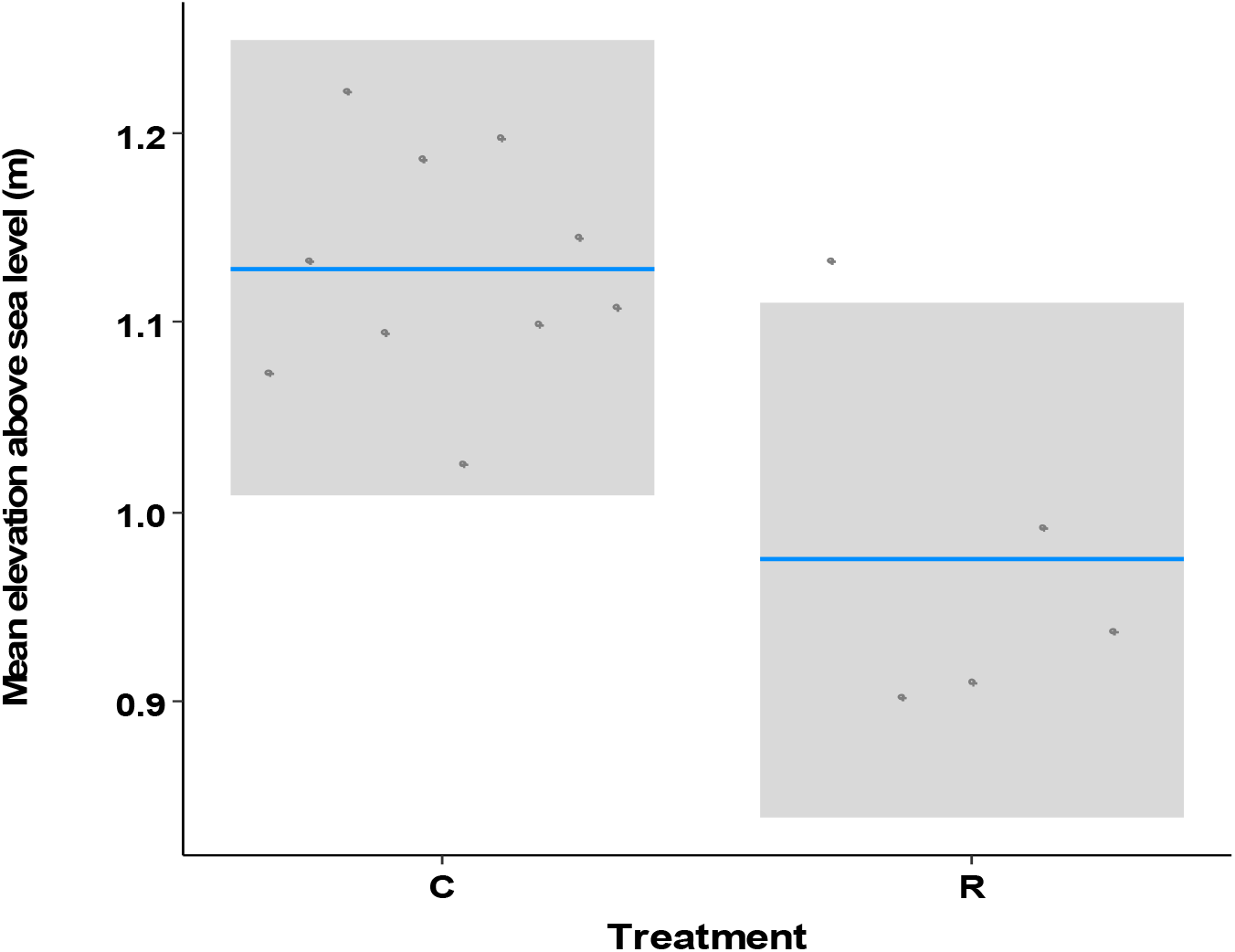
Effects of wrack on mean elevation above sea level considering data averaged across the final three timepoints. Plot shows the partial residuals from linear models accounting for experimental block as a function of treatment type (control vs removal). Significant effect detected (t = -3.19, p = 0.011) Elevation response derived from the average of three data collection time points (August, September and December 2019). Grey area represents the 95% confidence of the means.

**Fig. S7:**
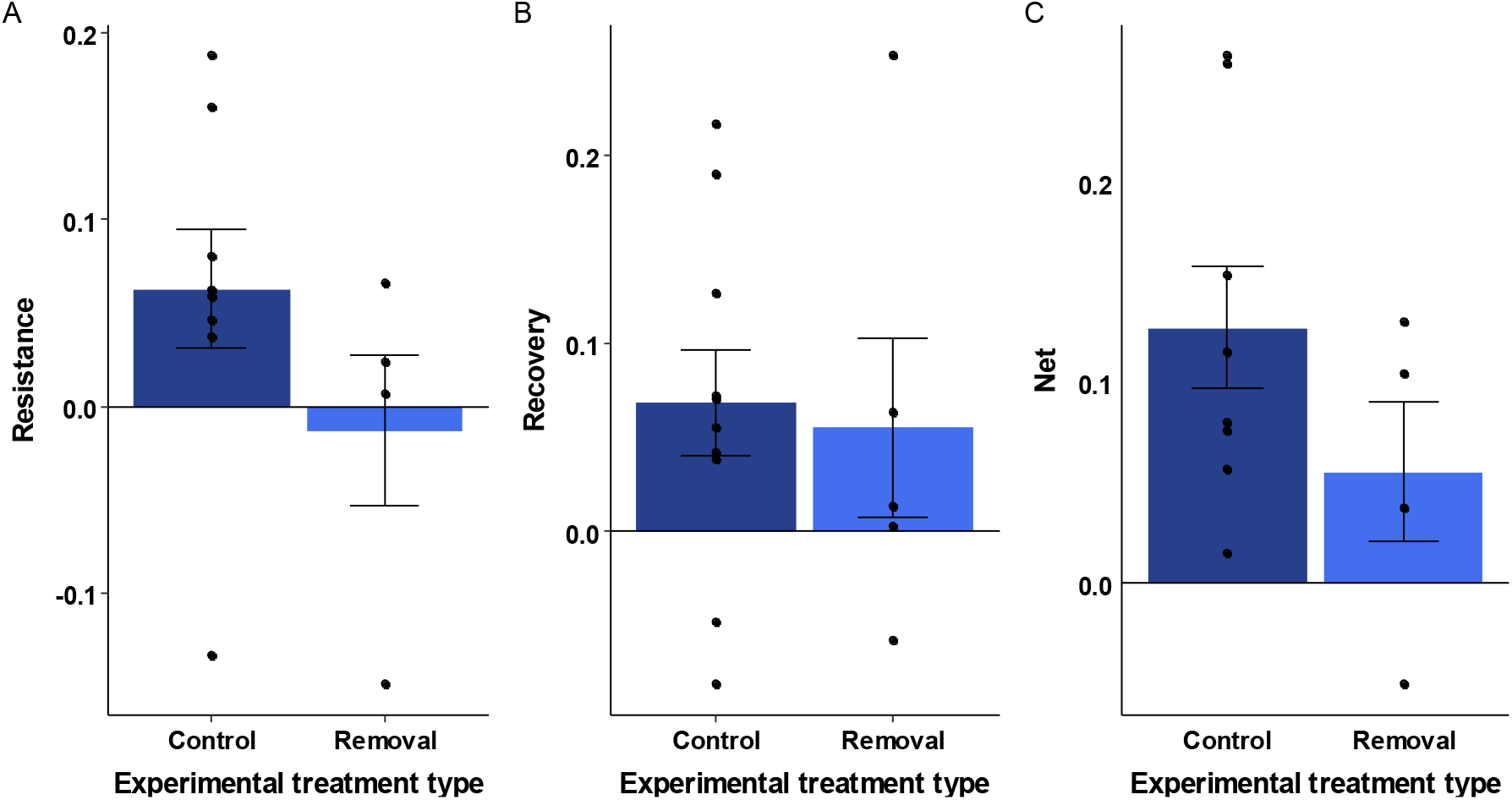
Effects of wrack removal on resistance metrics. All plots show treatment type (control vs removal) and standard error. Resistance metrics are: **A**) resistance - difference in mean elevation (m) above sea level between August and September 2019 (t = 2.103, p = 0.065), **B)** recovery - difference in mean elevation (m) above sea level before and after storm event (September and December 2019), (t = -0.295, p = 0.77) and **C)** net gain of mean elevation above sea level (total of resistance and recovery between August 2019 – December 2019), (t = -1.258, p = 0.24).

